# The role of the first transmembrane helix in bacterial YidC: Insights from the crystal structure and molecular dynamics simulations

**DOI:** 10.1101/2021.07.15.452481

**Authors:** Karol J. Nass, Ioana M. Ilie, Manfred J. Saller, Arnold J.M. Driessen, Amedeo Caflisch, Richard A. Kammerer, Xiaodan Li

## Abstract

The evolutionary conserved YidC is a unique dual-function protein that adopts insertase and chaperone conformations. The TM1 helix of *Escherichia coli* YidC mediates the interaction between the YidC chaperone and the Sec translocon. Here, we report the first crystal structure of *Thermotoga maritima* YidC (TmYidC) including the TM1 helix (PDB ID: 6Y86). The TM1 helix lies on the periplasmic side of the bilayer forming an angle of about 15 degrees with the membrane surface. Our functional studies suggest a role of TM1 for the species-specific interaction with the Sec translocon. The reconstitution data and the superimposition of TmYidC with known YidC structures suggest an active insertase conformation for YidC. Molecular dynamics (MD) simulations of TmYidC provide evidence that the TM1 helix acts as a membrane anchor for the YidC insertase and highlight the flexibility of the C1 region underlining its ability to switch between insertase and chaperone conformations. A structure-based model is proposed to rationalize how YidC performs the insertase and chaperone functions by re-positioning of the TM1 helix and the other structural elements.

## Introduction

YidC belongs to the YidC/Alb3/Oxa1 membrane protein family which is evolutionary conserved from archaea to man (1). The proteins of this family play an important role in the insertion and folding of membrane proteins in the bacterial, thylakoidal, mitochondrial inner membranes and the eukaryotic endoplasmic reticulum (2–4). Two conserved structural elements have been identified in all YidC proteins. First, the hydrophilic groove within the TMD (TM2-TM6), which represents the catalytic center of the protein. Second, the C1 region connecting TM2 and TM3, which consists of two helices (CH1 and CH2) connected by a short loop forming a hairpin-like structure (Fig. 1). Though sharing rather low sequence similarity, the hydrophilic groove and its ability to form lipid-mediated homo- or hetero-oligomers to promote the efficient protein translocation (2, 4) are conserved among the family members. The most studied protein from this family is YidC from *Escherichia coli* (EcYidC). Depletion of the YidC protein results in cell death due to the defective assembly of energy-transducing membrane complexes (5), this makes the YidC protein an attractive antibiotic target.

**Figure 1:**
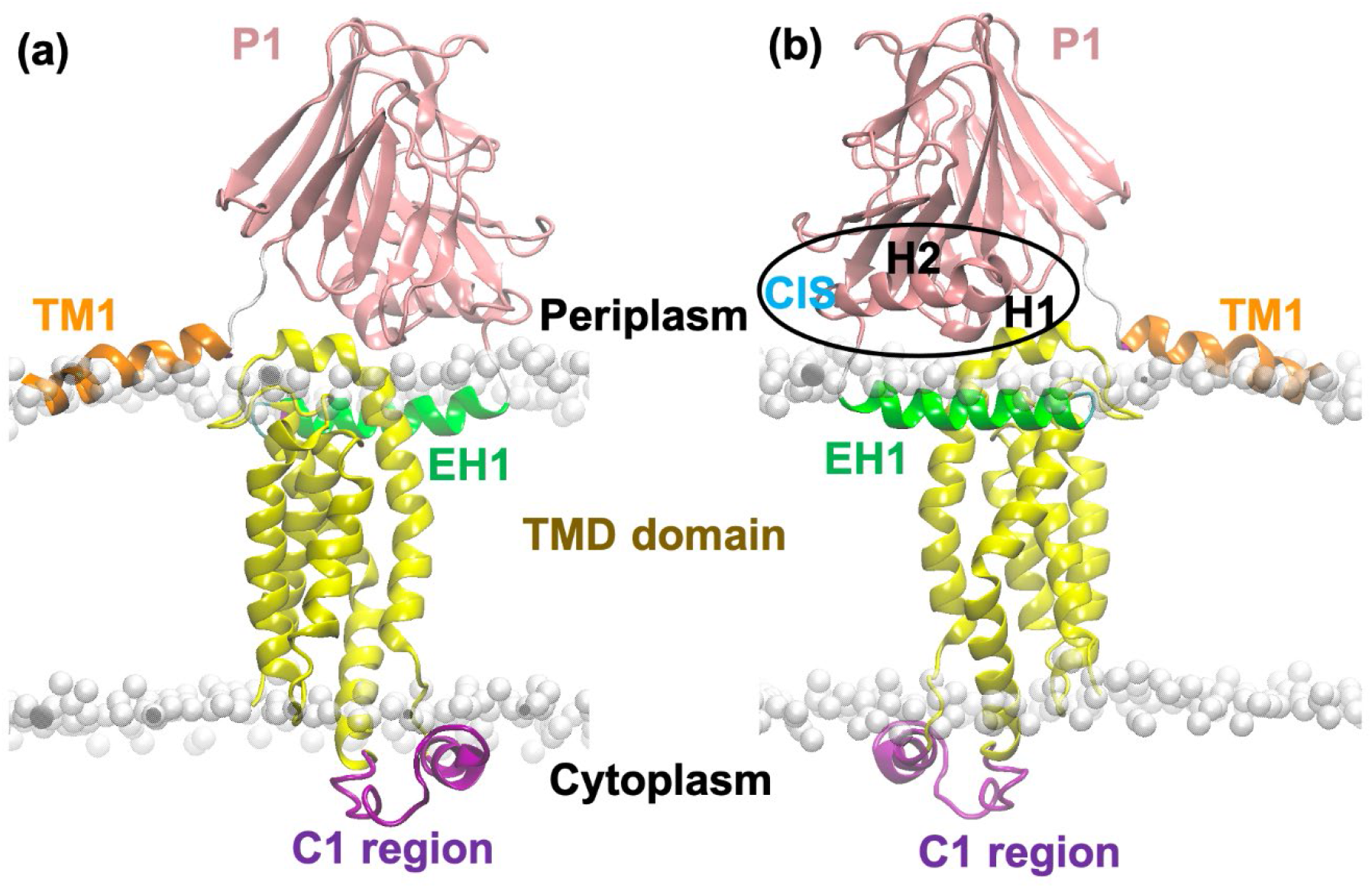
Overall structure of the TmYidC. (a) Front view of TmYidC. (b) Back view of TmYidC (PDB ID: 6Y86). TM1: orange (residues 1-19), P1: pink (23-225), EH1: green (226-244), TM region: Yellow (TM2 (248-280), TM3-TM6 (317-424)) and C1 region: magenta (282-308). A highly conserved interface structure (CIS) formed by packing C-terminal helix1 (H1) and helix2 (H2) against the β-supersandwich via hydrophobic interactions in P1 domain is shown in a black ellipse. The position of the membrane is shown in grey

The YidC chaperone is an integral part of the holo translocon (HTL), which folds and assembles multipass transmembrane proteins, as shown in the low-resolution cryo-EM structure of the SecYEG-SecDFYajC-YidC complex (6–8). The YidC chaperone facilitates the removal of non-mature membrane protein substrates from the lateral gate of the SecY protein, releasing the substrates into the lipid bilayer (9, 10). HTLs likely exist in a dynamic manner within the lipid bilayer for accommodating the diverse substrates and improving insertion efficiency (11, 12).

The hydrophilic groove is open towards the lipid bilayer and the cytoplasm but closed on the periplasmic side (13, 14). High B-factor values have been reported for the C1 region, indicating that this region fluctuates greatly near cytoplasmic bilayer surface, as also confirmed by the MD stimulations (13). The C1 region flexibility is required for the YidC insertase activity (15) and for YidC interactions with SecYEG (16). Notably, the YidC protein from Gram-negative bacteria contains an additional N-terminal TM1 helix and P1 domain between TM1 and TM2 (17). Functional experiments revealed a controversial role of TM1 helix for YidC activity. On the one hand, replacement of TM1 by the cleavable signal peptide of the maltose binding protein doesn’t affect the function of YidC (18). On the other hand, deletion of the TM1 helix compromises the insertion function of the YidC insertase, suggesting an anchor function of TM1 (19). Furthermore*, in vitro* site-directed and para-formaldehyde cross-linking experiments identified the TM1 helix of the EcYidC as a major contact site with the Sec translocon and the C1 region as an additional contact site for the Sec translocon. The low-resolution cryo-EM structure of the *E. coli* SecYEG-SecDFYajC-YidC HTL indicates that the TM1 helix intercalates at the lateral gate of the empty SecY and aligns the YidC hydrophilic groove to the lateral gate of SecY. Consequently, the hydrophilic groove of YidC and the lateral gate of SecY constitute a fusion protein-conducting channel with unique properties (10, 20). The fusion protein-conducting channel has a smaller pore size with less hydrophobicity and higher anion selectivity in comparison to the SecY channel alone. Furthermore, the fusion protein-conducting channel lowers the energy barrier for substrate releasing to the lipid bilayer (8, 21–25).

Here, we address the structural and functional roles of the TM1 helix. By functional studies with chimeric proteins of EcYidC and TmYidC, we provide evidence that the TM1 segment is important for the species-specific interaction with SecY. Furthermore, we present a TmYidC crystal structure in which the TM1 helix is visible for the first time (PDB ID: 6Y86). Molecular dynamics simulations reveal that the angle between the axis of the TM1 helix and the membrane surface is about 15 degrees and that the N-terminus of the TM1 helix is embedded in the membrane acting as an anchor. The superimposition of TmYidC with the known YidC structures yields a root mean square deviation of 3.3 Å from the active structure of the gram-positive *Bacillus halodurans* YidC insertase (BhYidC), suggesting an active TmYidC insertase, consistent with our reconstitution data. Our combined experimental and computational study shows that the TM1 helix, the EH1 helix and the C1 region of YidC protein can adopt different conformations for insertion and interaction with the Sec translocon. We propose a structure-based model characterizing the insertase and chaperone conformations of YidC protein.

## Results

### Functional analysis of TM1 from *Thermotoga maritima* YidC

Based on crosslinking data, we propose that the TM1 helix of YidC needs to interact with SecY to allow YidC to perform its function as a chaperone, thereby suggesting a species-specific functional role of the TM1 helix. To test this hypothesis, EcYidC, TmYidC and a TmYidC chimera containing the TM1 helix of EcYidC (YidC61) (Fig.2a) were expressed in *E. coli* FTL10 (see Supplementary Information Table 1). In this strain, the *yidC* gene is under the control of the *araBAD* promotor. Since Ec_YidC is essential for cell viability, the growth of *E. coli* FTL10 is dependent on the presence of arabinose in the medium (26). For the complementation assays, plasmids pTrcTmaHis and pTrce61-t396His (see Supplementary Information Table 2) were introduced into the FTL10 strain and cells were grown with glucose instead. In YidC61, the TM1 helix (N-terminal 47 amino acids) of TmYidC was replaced with the N-terminal (61 residues) of EcYidC. The topology of these proteins is shown in Fig. 2a. In addition, the EcYidC (pTrcyidc) and the empty vector (pTrc99A) were used as positive and negative controls, respectively (see Supplementary Information Table 2). Transformed cells were tested for growth in liquid Luria-Bertani (LB) medium. As expected, all cells were viable in the LB medium supplemented with arabinose (Fig. 2b, upper panel). However, under conditions, which repress the expression of chromosomal *yidC* (glucose, IPTG), EcYidC (positive control) and YidC61 complemented the *E. coli* FTL10 growth defect (Fig. 2b, lower panel). Cells transformed with the empty vector and TmYidC showed poor growth under these conditions. One of the possible reasons for the lack of YidC complementation by TmYidC is that the TM1 of TmYidC is not able to interact with *E. coli* SecY (EcSecY) to form a HTL, which can insert and fold the substrates more efficiently and is required for growth. Western blot experiments using purified EcSecY, EcYidC and TmYidC confirm the existence of a weak and transient interaction between EcSecY and endogenous EcYidC, but not when TmYidC is overexpressed (see Supplementary Information Fig.1). TmYidC can be expressed and purified from *E. coil* membrane suggesting that no complementation is not due to a lack of protein expression and/or a failure in membrane targeting. These data underscore the importance of the TM1 helix for YidC activity and might reflect a species-specific function, for the interaction with SecY (10). An alignment of different YidC proteins shows that the TM1 helix of all gram-negative bacteria YidC share some similarities, yet the isoelectric point and the net charge of TM1 of EcYidC and TmYidC are different*, i.e.,* 4.3 with one negatively charged residue and 8.5 with one positively charged residue, respectively (see Supplementary Information Fig.2).

**Figure 2:**
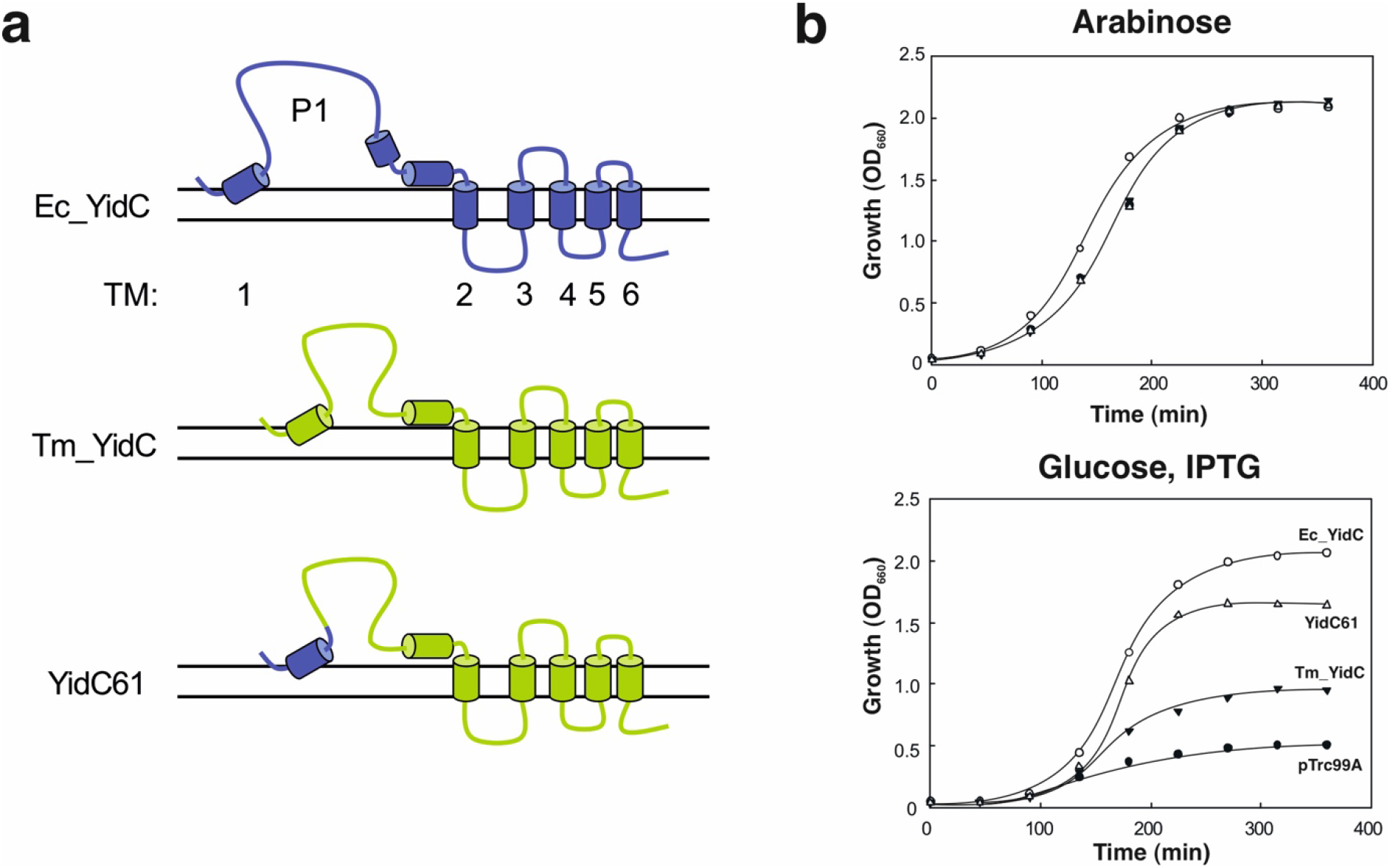
Complementation of the growth defect of an *E. coli* YidC depletion strain by chimeric *E. coli* and *T. maritima* YidC proteins. (a) Topology models of the EcYidC (blue) and the TmYidC (green) proteins. The α-helical regions (TM and hydrophilic helices H1 and H2) are represented by cylinders, solid lines indicate soluble terminal and loop regions. YidC61 is a chimeric protein consisting of the TM1 from EcYidC (N-terminal, blue) and TmYidC (C-terminal, green). (b) The *E. coli* FTL10 cells were grown in a medium supplemented with either 0.5% (wt/vol) arabinose (top panel) or glucose and 50 μM IPTG (bottom panel). Cells containing the empty plasmid pTrc99A (filled circles) or plasmids for the expression of EcYidC (open circles), YidC61 (open triangles) and TmYidC (filled triangles) were grown under induction and repression conditions.

### The crystal structure of TmYidC and molecular dynamics simulations suggest that the TM1 helix is a membrane anchor for the YidC insertase

We crystallized TmYidC using the vapor diffusion method in presence of the detergent dodecyl maltoside (DDM) and solved the structure by molecular replacement using the crystal structure of the P1 domain of TmYidC as search model (unpublished). The final model was refined to 3.4 Å resolution (Table 1). The asymmetric unit contains one TmYidC molecule. This data is consistent with size exclusion chromatography experiments showing TmYidC existing as a monomer in solution containing DDM (see Supplementary Information Fig. 3). Structure analysis of TmYidC was carried out using published YidC structures (13, 14, 27–30).

**Table 1.**
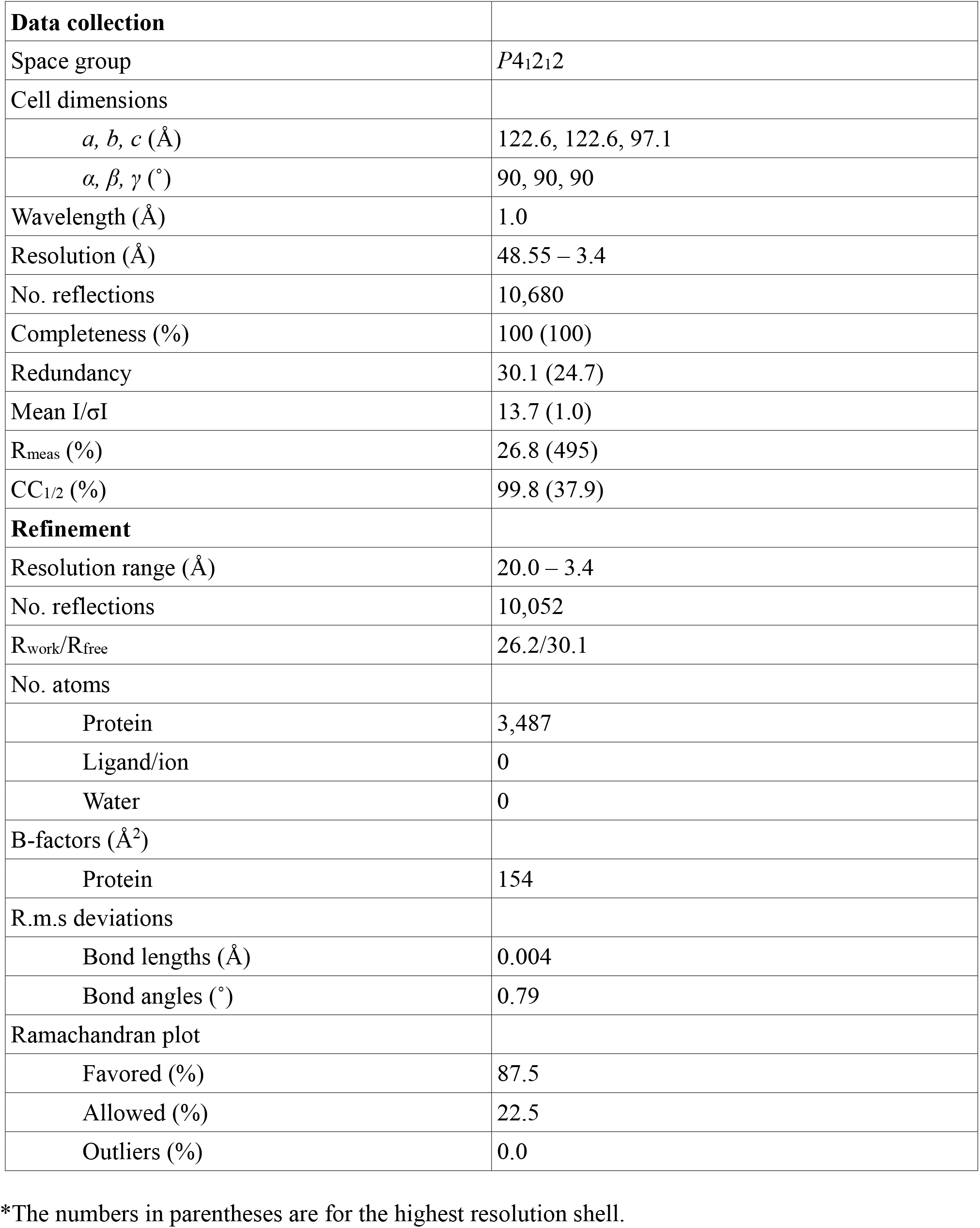
Data collection and refinement statistics

The electron density of TmYidC is well defined for residues 1-425 including the TM1 helix (residues 1-22). Residues 426-445 are not visible in the structure, indicating the flexibility of the C-terminus in the absence of the ribosome. Front and back view of the TmYidC structure are shown in Fig. 1a and Fig. 1b, respectively. The overall structure of TmYidC is well defined with all structural elements visible (PDB ID: 6Y86). The TM1 helix lays on the periplasmic membrane face, forming a roughly 15-degree angle with the membrane and is located on the opposite side of the entrance to the hydrophilic groove. To further investigate the orientation of the TM1 helix we performed molecular dynamics simulations starting from the crystal structure of TmYidC, which was first embedded in a POPE:POPG (3:1 ratio) bilayer to mimic the membrane environment of the YidC insertase. Ten independent simulations were carried out for a total sampling of 4 μs. The results reveal that the tilt angle of the TM1 helix with respect to the membrane surface (Fig. 3a) fluctuates in the [-10, +30] degrees interval (Fig. 3b) with an average of 13 degrees and a standard deviation of ±8 degrees. The small standard error of the mean (±3.4 degrees, evaluated as the standard deviation of the ten average values along the individual runs) suggests that the sampling has reached convergence. Secondary structure analysis shows that TM1 helix preserves its α-helical structure throughout the simulations except for the C-terminal residues Leu18-Phe19, which sample mainly coil- and bend-like conformations (Fig. 3c).

**Figure 3:**
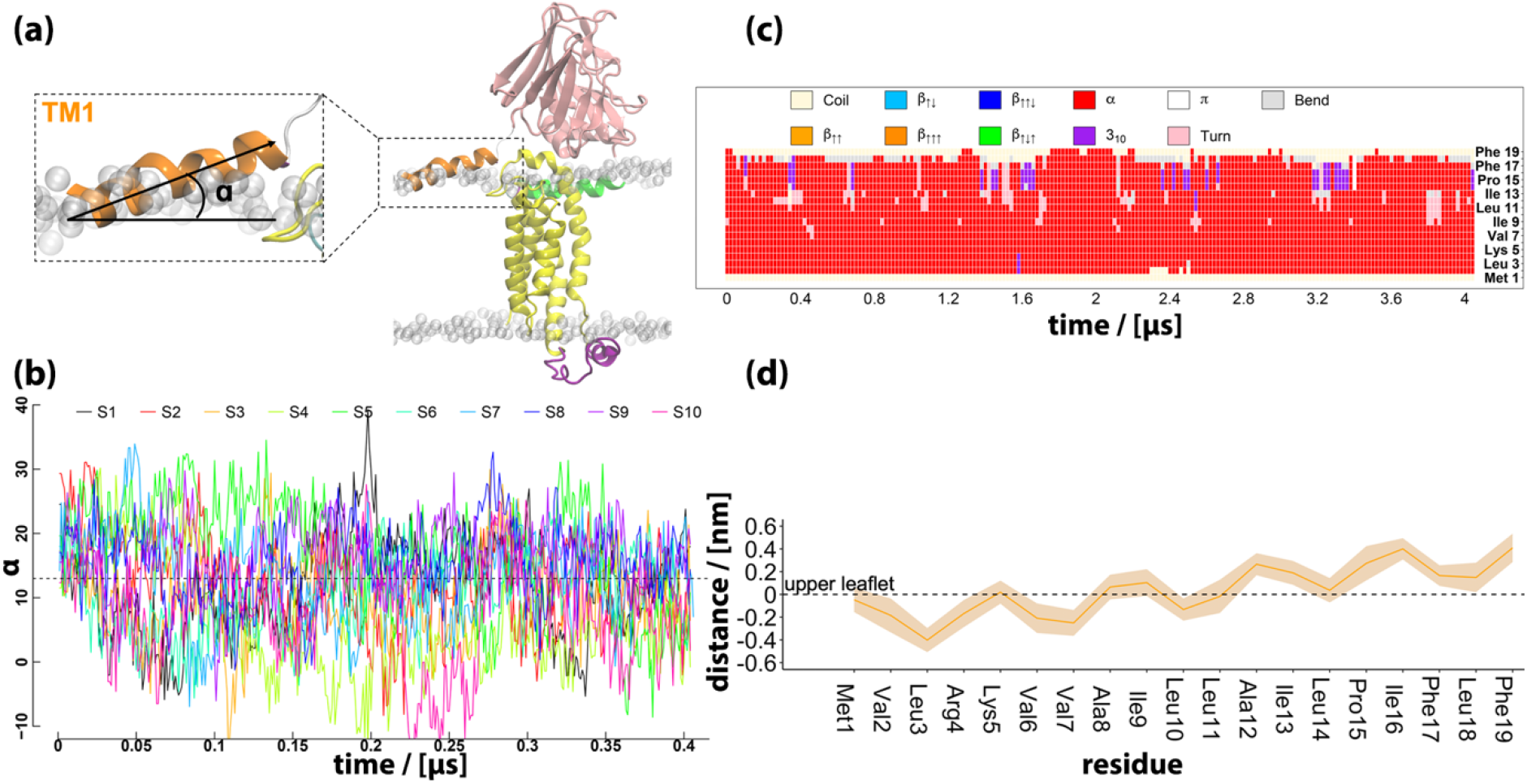
Molecular dynamics characterization of the TM1 helix (residues 1-19). (a) Definition of the tilt angle between the TM1 helix and the membrane surface. The angle α was calculated between a vector connecting the centers of mass of the backbone atoms of residues 2-5 and 14-17 (N-terminal turn and C-terminal turn of the TM1 helix, respectively) and the membrane plane. (b) Time series of the angle α. The horizontal dashed line represents the average over the 10 simulations. (c) Time series of the sequence profile of the secondary structure. The figure shows the cumulative sampling of 10 μs. (d) The embedding of the TM1 helix in the membrane is illustrated by the sequence profile of the distance between the Cα atoms of the TM1 helix and the average position of the phosphorus atoms of the phospholipids in the upper leaflet (black dashed line). The orange area represents the standard error of the mean calculated as the standard deviation of the 10 average values over the 10 individual runs.

In the crystal structure, the TM1 helix is tilted on the membrane surface and makes crystal contacts with two symmetry related molecules in the crystal packing. In this conformation, TM1 could fulfill its role as an anchor for localizing the YidC protein into the lipid bilayer, which is in full agreement with deletion experiments (19) and the complementation experiment shown above. The molecular dynamics simulations of TmYidC in the lipid bilayer demonstrate that the N-terminus of the TM1 helix is embedded in the membrane (Fig. 3d), thereby reinforcing the anchor role of TM1.

### The P1 domain and the EH1 helix of YidC can switch between insertase and chaperone conformations

The P1 domain of YidC forms a compact and rigid structure consisting mainly of a β-sandwich, which accounts for more than half of the total amino acids of the protein (31), and is involved in the interaction with SecF (32). The P1 domain of EcYidC is oriented towards the periplasmic membrane through its conserved C-terminal region together with the amphipathic EH1 helix (13, 31). While deletion of the β-sandwich of the P1 domain does not abolish the function of EcYidC (19), deletion of the C-terminal region (311-327) composed of two short α-helices (helix 1 and helix 2) impairs cell viability and membrane insertion of a number of Sec-dependent substrates (31, 32). These two short α-helices form a highly conserved interface structure (CIS), which packs against the β-sandwich (31). The number and the arrangement of β-sheets in the P1 domain of the EcYidC is different from that of TmYidC. Accordingly, we did not use the P1 domain of EcYidC as a template for solving the structure of TmYidC by molecular replacement (31, 33). The P1 domain of TmYidC (23-224) consists of fifteen β-sheets, one 3/10 helix and one CIS, while the EcYidC P1 domain is composed of nineteen β-sheets, two 3/10 helices and one CIS. As the CIS domain and the EH1 helix contribute to the orientation of the P1 domain to the membrane (31), we used CIS as hallmark for the P1 domain orientation/conformation of the YidC protein. The position of CIS in the EcYidC insertase (PDB ID: 6AL2), is different from the EcYidC chaperone within the HTL (PDB ID: 5MG3) (Fig. 4a). The CIS of the YidC insertase is closer to the periplasmic membrane plane compared to that of the YidC chaperone within the HTL. Interestingly, helix 2 of CIS is invisible in the EcYidC chaperone within the HTL. Although the position of CIS of TmYidC is similar to that of the EcYidC insertase (Fig. 4b), helix 1 (199-205) and helix 2 (210-217) of the TmYidC do not overlay with helix 1 and helix 2 of the EcYidC (Fig. 4c). Moreover, helix 2 of TmYidC is parallel to the EH1 helix, and helix 1 forms approximately a 40-degree angle with respect to helix 2. Such an arrangement pushes the EH1 helix closer to the SecY protein (Fig. 4b and 4c), indicating the coupling of the P1 domain to the TM domain through the EH1 helix, which consequently modulates the interaction of YidC with SecY.

**Figure 4:**
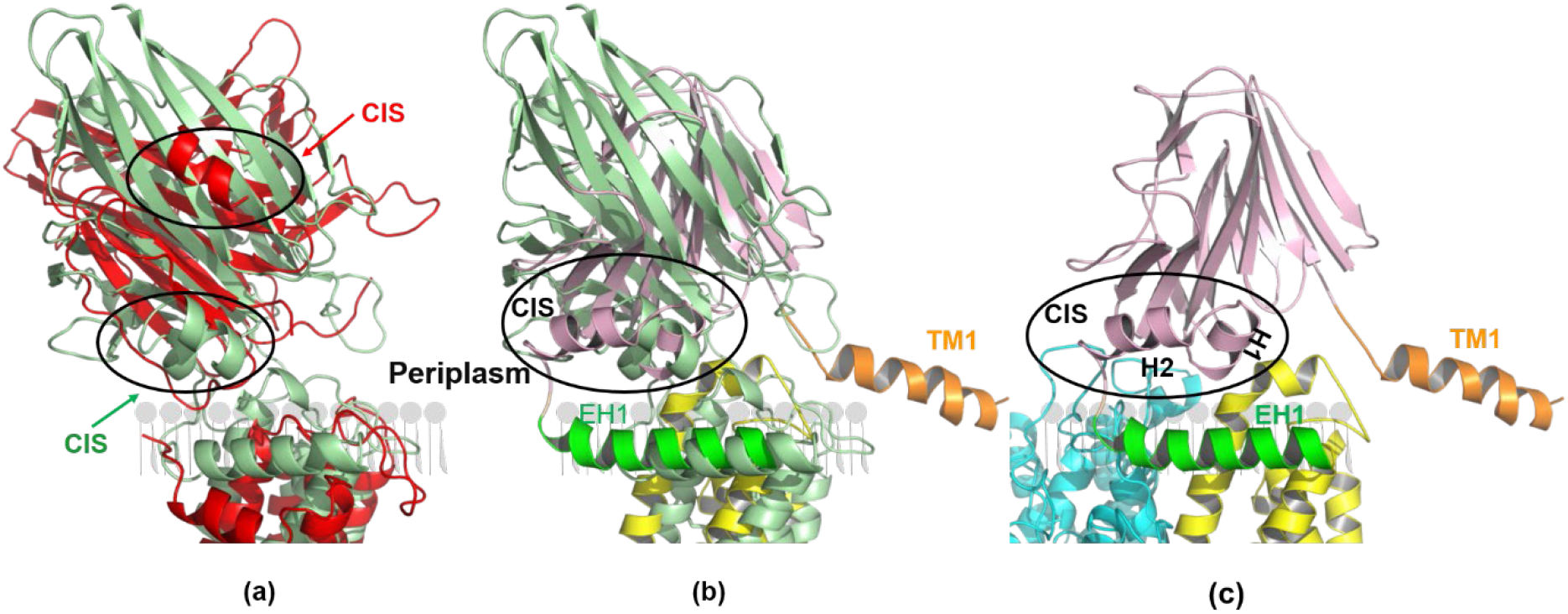
Conformations of the P1 domain. (a) Comparison of the P1 domains of the EcYidC insertase (PDB ID: 6AL2, pale green) and the EcYidC chaperone (PDB ID: 5MG3, chain C, red). The CIS of region of both structures is marked black ellipses arrowed in pale green and red respectively. (b) Comparison of the P1 domain of the TmYidC (PDB ID: 6Y86, pink) and the EcYidC insertase (PDB ID: 6AL2, pale green). The CISs of both proteins are marked in black ellipse. (c) Position of CIS with helix 1 (H1) and helix 2 (H2) of our structure and SecY (PDB ID: 5MG3, chain Y, cyan). The structures were aligned by structural overlap of the common YidC domains.

The position and the length of the EH1 helix are important for its function as a mechanical lever to coordinate the movement of the YidC protein for releasing the nascent chain from the hydrophilic groove into the lipid bilayer (34). The molecular dynamics simulations show that the EH1 helix of TmYidC is embedded in the lipid bilayer and aligned in a parallel fashion to the membrane surface (Fig. 5). A similar parallel arrangement, yet on the surface of the membrane, is recorded by the 11 amino acid long EH1 helix (340-351) of EcYidC insertase (PDB ID: 6AL2). On the other hand, the EH1 helix of EcYidC chaperone within the HTL (PDB ID: 5MG3) is inserted into the lipid bilayer and is 8 amino acids long (339-347) (Fig. 6a). The arrangement of the 18 residues of the EH1 helix of TmYidC (residues 226-244) overlap well to those of EcYidC insertase (Fig. 6b) and BhYidC insertase (29-56) (Fig. 6c) indicating an insertase conformation of the EH1 helix.

**Figure 5:**
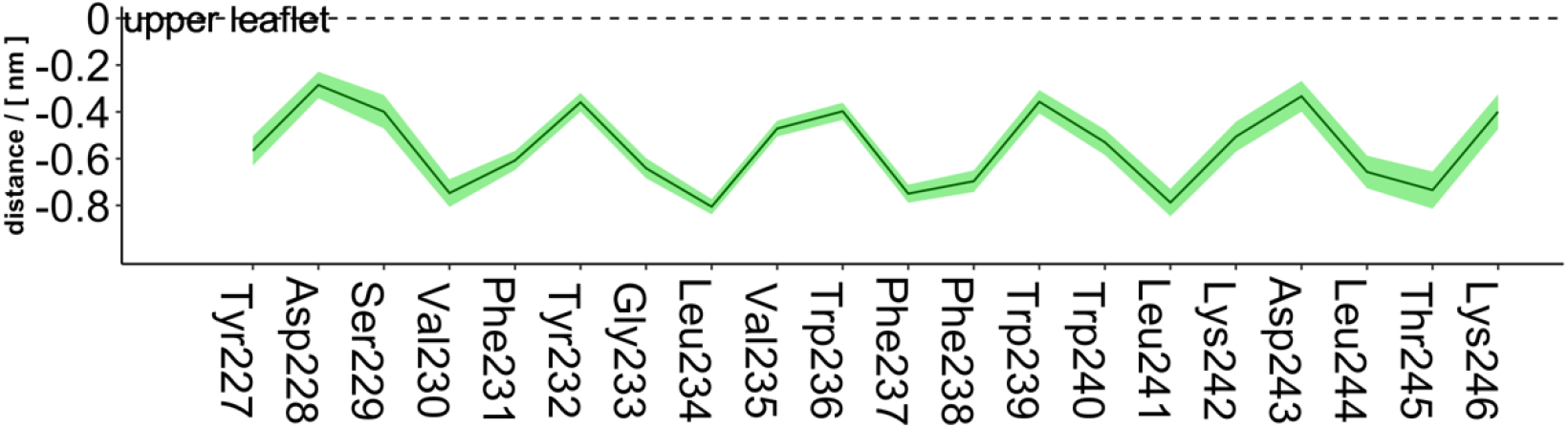
Embedding of the EH1 helix in the membrane. The embedding is illustrated by the sequence profile of the distance between the C_α_ atoms of the TM1 helix and the average position of the phosphorus atoms of the phospholipids in the upper leaflet (black line). The green area represents the standard error of the mean calculated as the standard deviation of the 10 average values over the 10 individual runs.

**Figure 6:**
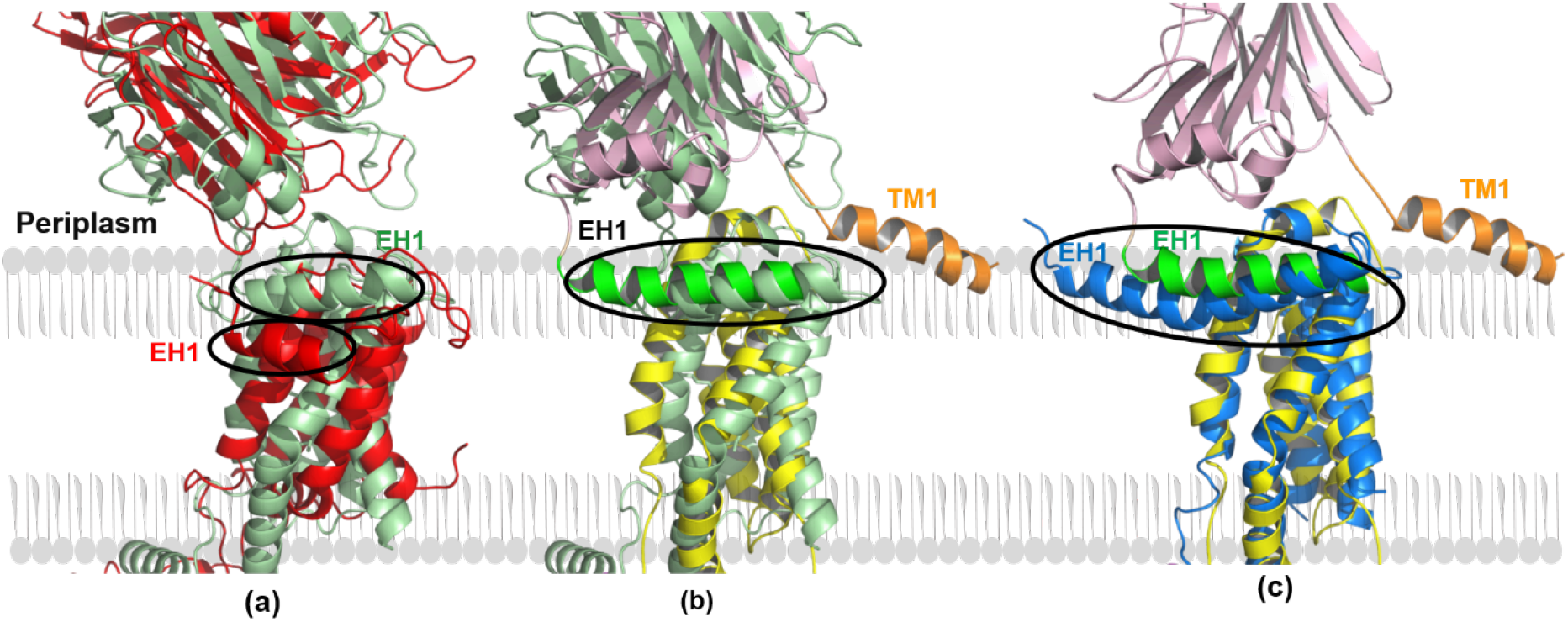
Conformations of the EH1 helix. (a) Comparison of the EH1 helix of the EcYidC_insertase (PDB ID: 6AL2, pale green) and the EcYidC_chaperone (PDB ID: 5MG3, chain C, red). The EH1 helix of both structures is marked with black ellipses marked in pale green and red respectively. (b) Comparison of the EH1 helix of the TmYidC (PDB ID: 6Y86, green) and EcYidC_insertase (PDB ID: 6AL2, pale green) marked in black ellipse. (c) Comparison of the EH1 helix of the TmYidC (green) and the BhYidC_insertase (blue) (PDB ID: 3WO6) marked in black ellipse.

### The C1 region of TmYidC adopts an insertase conformation

The electron density of the C1 region of TmYidC (residues 282-308) is well defined with a high B-factor, indicating the flexibility of C1 region is indeed a general characteristic of the YidC protein. This result is supported by molecular dynamics simulations, which show that the C1 region records deviations up to 2.5 nm from the crystal structure (Fig. 7a). Of the two helices solved in the crystal structure (Fig. 1), only one (residues 299-308) preserves its helical conformation, whereas the second samples mainly coil and turn rich conformations (Fig. 7b). The flexibility of the C1 region in the lipid bilayer environment is shared by BhYidC insertase, as shown in previous simulations (13). Furthermore, the structural overlap of the crystal structures of the EcYidC insertase and the EcYidC chaperone within the HTL displays evident differences in the arrangements of the CH1 and CH2 helices of the C1 region (Fig. 8a). Additionally, further comparisons of the C1 region reveal structural similarities to EcYidC and TmYidC (Fig. 8b) and structural differences to the EcYidC chaperone in the presence of SecY (Fig. 8c). For the chaperone, the C1 region of EcYidC within the HTL is in contact with the SecY, while the C1 region in the insertase conformation would not interact with the SecY protein (13, 29). The diverse arrangements of the C1 region in the crystal structures and the flexibility of this domain as highlighted by the molecular dynamics simulations (Movie S1) underline its ability to switch between insertase and chaperone conformations.

**Figure 7:**
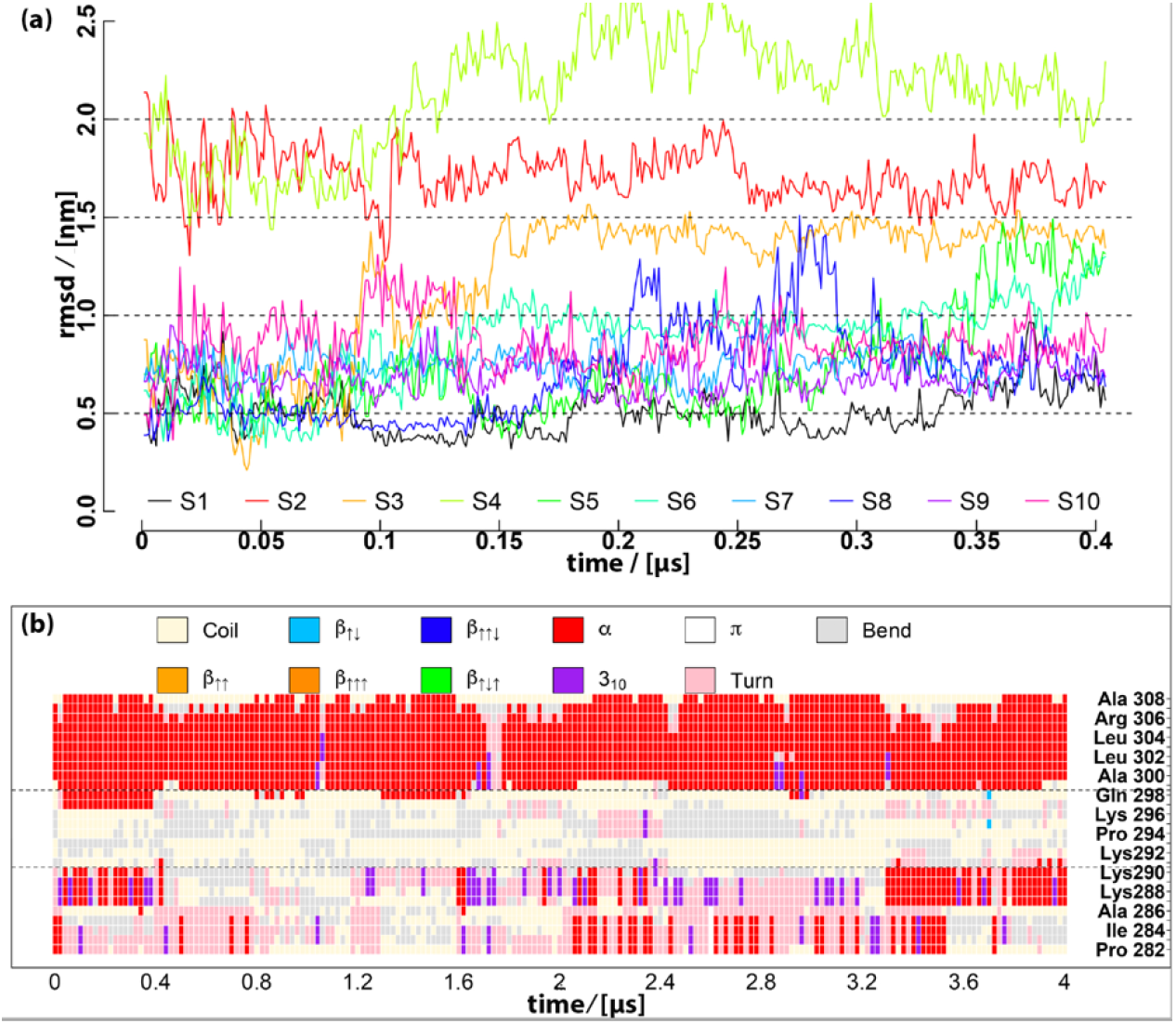
Characterization of the C1 domain (residues 282-308). (a) Root mean square deviation of the Cα-atoms in the C1 domain from the crystal structure (PDB ID: 6Y86). The alignment was done on the Cα-atoms of the TM2-TM6 domain excluding the C1 region. (b) Secondary structure assignment of the residues in C1. The horizontal dashed lines separate the two helices in the crystal structure and the connecting loop.

**Figure 8:**
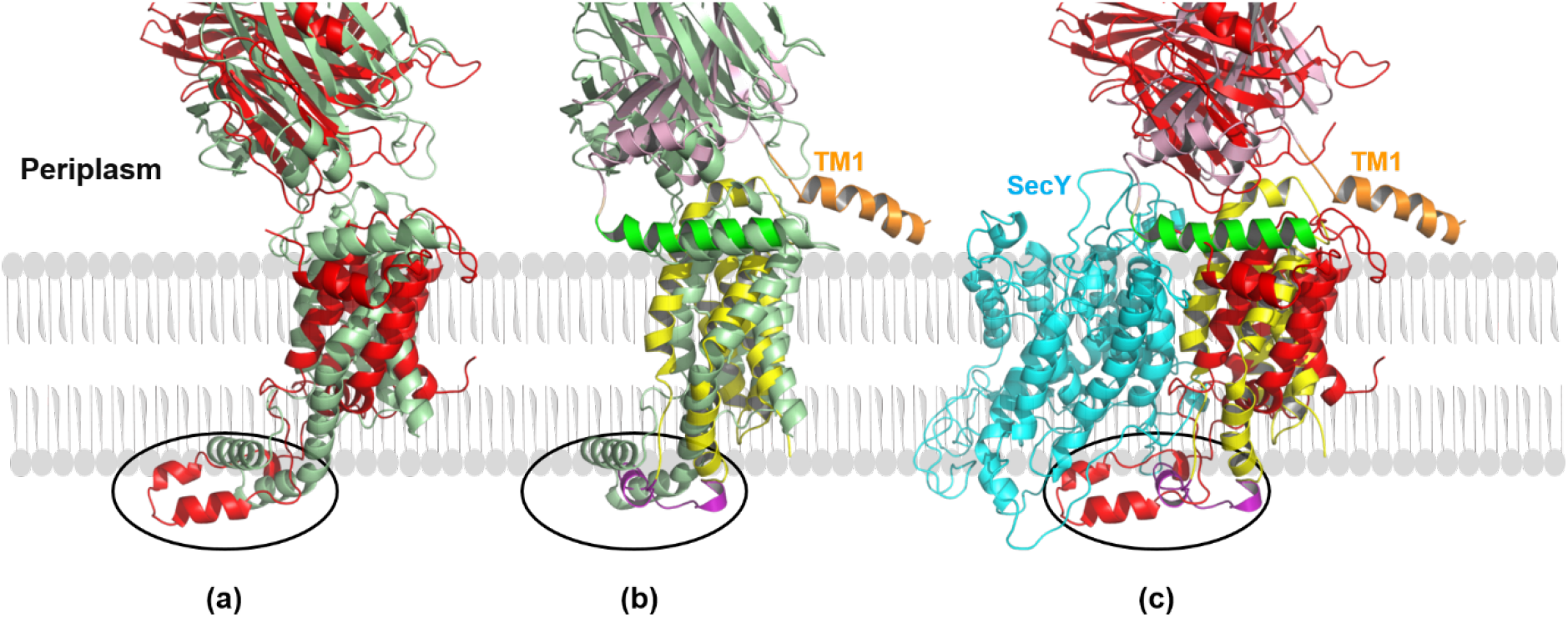
Conformations of the C1 region (marked in the black ellipses). (a) Comparison of the C1 region of the EcYidC insertase (PDB ID: 6AL2, pale green) and the EcYidC chaperone (PDB ID: 5MG3, chain C, red). (b) Comparison of the C1 region of the TmYidC (PDB ID: 6Y86, magenta) and the EcYidC insertase (PDB ID: 6AL2, pale green). (c) Comparison of the C1 region of the TmYidC (magenta) and the EcYidC chaperone (PDB ID: 5MG3, chain C, red) in the presence of the SecY protein (PDB ID: 5MG3, chain Y, cyan). The structures were aligned by structural overlap of the common YidC domains.

### The hydrophilic groove of TmYidC represents an active insertase conformation

The hydrophilic groove of TmYidC is located within TM2-TM6 and opens to the membrane and the cytoplasmic side (Fig. 1a). The structure of the hydrophilic groove is conserved within the YidC/Oxa1/Alb3 family members. Comparison with all known YidC insertase structures reveals that each TM helix of the hydrophilic groove of TmYidC superimposes to the TM helices of the BhYidC (PDB ID: 3WO6) with a RMSD of 3.3 Å (Fig. 9a), and to the EcYidC (PDB ID: 6AL2) with a RMSD of 9.7 Å (Fig. 9b). The BhYidC (PDB ID: 3WO6) has been identified as an active insertase (13), suggesting that TmYidC insertase is likely in an active conformation. This is in good agreement with reconstituted TmYidC mediating the membrane insertion of the *in vitro* synthesized c-subunit of *E. coli* ATPase (see Supplementary Information Fig. 4).

**Figure 9:**
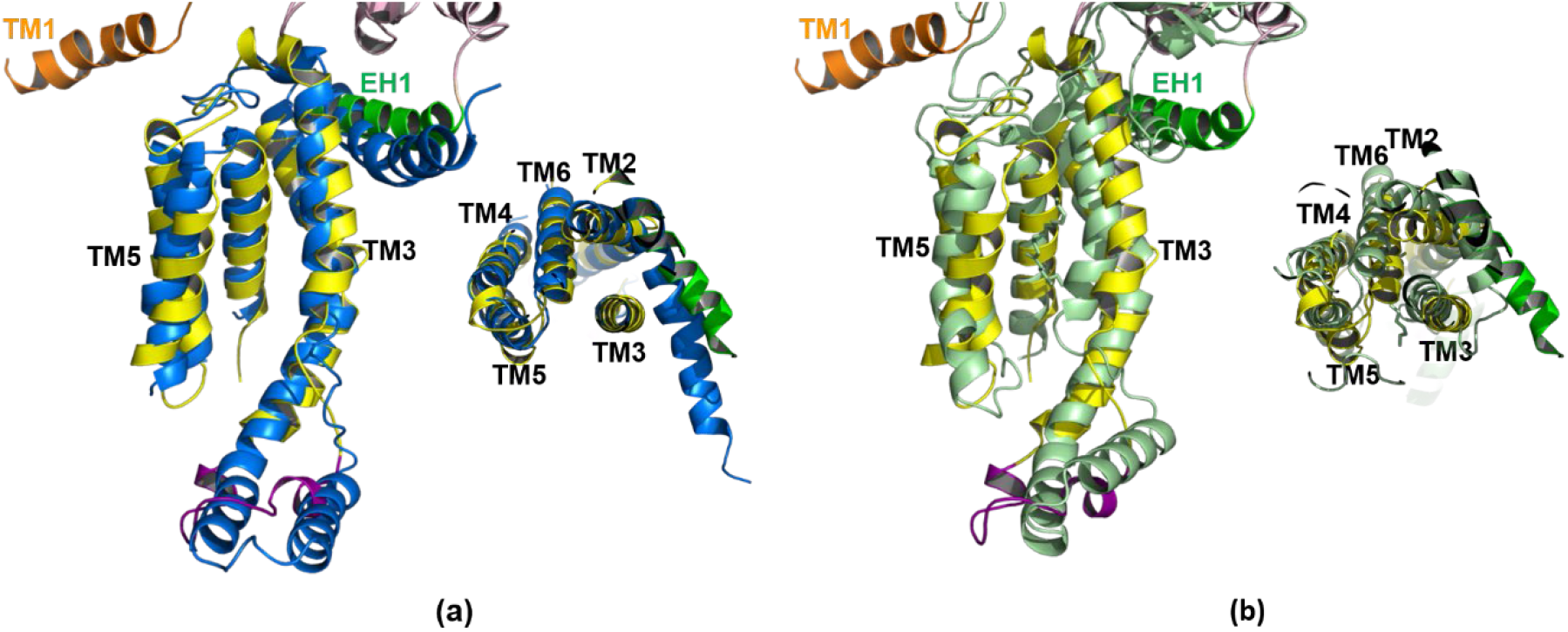
Comparison of the TM region of TmYidC with BhYidC insertase (PDB ID: 3WO6) and EcYidC insertase (PDB ID: 6AL2). (a) Comparison of the TM region of TmYidC (PDB ID: 6Y86, residues 245-425, yellow) with BhYidC Insertase (PDB ID: 3WO6, blue), side and top views. For clarity, TM1 (Orange) and EH1 (Green) of TmYidC are also shown. (b) Comparison of the TM region of TmYidC (PDB ID: 6Y86, residues 245-425, yellow) with EcYidC insertase (PDB ID: 6AL2, pale green), side and top views.

### A structural model explaining the dual function of the YidC protein

The structural analysis of TmYidC demonstrates that the YidC protein has two fundamental structural modules. The first/basic module is a catalytical unit, *i.e.,* the conserved hydrophilic groove required for its insertion activity. The second is the interacting module, comprising TM1, the P1 domain, the EH1 helix and the C1 region required for the interaction with the Sec protein. The interacting module can adapt between insertase and chaperone conformations. Here, we propose a plausible structural model to explain the insertase and chaperone conformations.

In the insertase conformation, TM1 is tilted above the periplasmic membrane, the EH1 helix is embedded in the lipid bilayer at the periplasmic side arranged in a parallel fashion to the membrane surface, and the C1 region can be in the proximity of the TM3 helix of the SecY protein (10) (Fig. 10a). In the chaperone conformation, based on the cross-link study (10), we propose that TM1 lays between TM3 and TM5 acting as the major contacting site with SecY, the EH1 helix is tilted and inserted into the membrane and pushed more closely to SecY, and the C1 region is contacting TM1 and TM10 of SecY (Fig. 10b). This model explains how the YidC protein can function both as an insertase and as a chaperone by rearranging its interacting modules, including TM1 in Gram-negative bacteria. For the YidC/Oxa1/Alb3 family members without the TM1, the EH1 and the C1 region could function similarly. The basic module of the YidC/Oxa1/Alb3 family of proteins can perform the insertion function with low efficiency. The interacting module of the YidC protein incorporating Sec proteins enables the insertion of diverse protein substrates and increased insertion efficiency.

**Figure 10:**
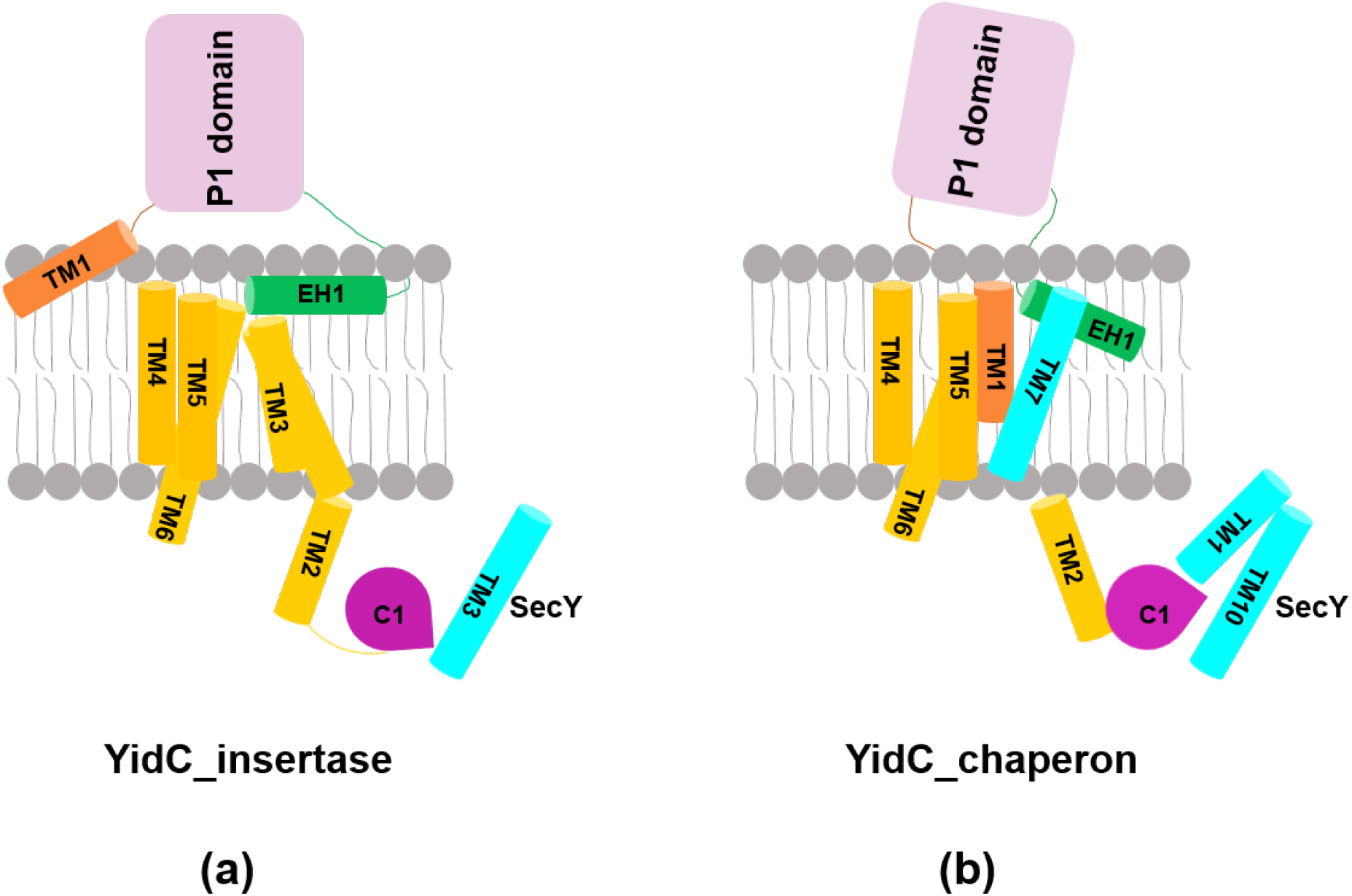
Structural models of the YidC insertase (a) and the YidC chaperone (b) functions of Gram-negative bacteria. For clarity, only selected helices of the SecY protein are shown (cyan).

## Discussions and Conclusions

We present here a 3.4 Å-resolution structure of an active YidC insertase with the TM1 helix resolved (PDB ID: 6Y86). By maintaining the stability and functionality of the hydrophilic groove in the TM region for the insertase activity, YidC deploys its chaperone function by interacting with Sec proteins using the TM1 helix, the P1 domain, the EH1 helix and the C1-region (10, 35). The structure solved here encompasses the TM1 helix and shows for the first time how the TM1 helix can act as a membrane anchor in the insertase conformation. Furthermore, it provides the structural basis for cross-linking experiments demonstrating that the TM1 helix is the major contacting site in the presence of SecY (10). The presence of both TM1 and C1 in TmYidC suggests that these structural elements of the YidC protein from *T. maritima* are less flexible than those of the other YidC insertases in presence of the detergent dodecyl-maltoside (DDM). The previously published TmYidC structure did not contain the TM1 helix and the C1 region, which might be due to using a shorter C10 alkyl chain of decyl-maltoside (DM) instead of a longer C12 DDM detergent, DM is harsher than DDM, which might disturbing these regions (36). The amphipathic EH1 helix connecting the P1 and TMD domains facilitates the coordinated movements of these domains to trigger the release of nascent chains into the membrane (34).

TmYidC is crystallized in the P 41 21 2 space group. In this lattice, the TM1 helix is stabilized by crystal contacts with two symmetry related molecules in the crystal lattice. The highly hydrophobic TM1 can lie in the plane of the bilayer or insert into the lipid bilayer. During YidC biogenesis, the TM1 helix has to be inserted into the lipid bilayer at certain points during synthesis. Whether transmembrane or surface bound, the TM1 helix could induce changes in the orientation of the P1domain and possibly the TMD domain. Our MD stimulations of TmYidC embedded in a POPE:POPG bilayer show that TM1 forms an angle of about 15 degrees with the membrane surface consistent with the crystallographic observation. Furthermore, the N-terminus of the TM1 helix is embedded in the membrane. The consequences of the tilted TM1 helix of TmYidC are twofold. First, could induce an insertase-compatible P1 domain and EH1 helix. Second, it could make the interaction with SecY more easily. The TM1 helix is therefore a definitory structural element, which can enable the dual-function of the YidC protein.

Combined with structural analysis of available YidC structures and molecular dynamics simulations, we propose a model that rationalizes how YidC can perform its dual function as an insertase or chaperone. To validate this model, structure determination of a high-resolution YidC-SecYEG complex will be required.

## Materials and Methods

### Strains and plasmids

#### Expression and Purification of YidC of *Thermotoga maritima*

##### Crystallization and Data collection

See Supplementary Information

##### Structure determination and refinement

The structure of TmYidC was determined to 3.4 Å by molecular replacement with the crystal structure of the P1 domain of TmYidC (unpublished) using PHASER (37). In resulting maps, additional electron density was observed, presumably for the TM domain of TmYidC. However, the electron density map was not sufficiently interpretable to build the TM domain. To improve initial phases, molecular replacement was performed using the P1 domain and TM domains of TmYidC determined separately (unpublished) as search models with the C1 region omitted. The model with the P1 domain and the TM region was placed into the additional electron density observed before and it was further rebuilt manually using COOT (38) and refined in Phenix (39). Data collection and refinement statistics are summarized in Table 1. The atomic coordinates and structure factors were deposited in the Protein Data Bank, under the accession code 6Y86.

### Molecular dynamics simulations

#### System preparation – YidC structure

The crystal structure of TmYidC (PDB ID: 6Y86) was aligned along the membrane normal using the Orientations of Proteins in Membranes (OPM) database (40) and subsequently embedded in a 75% 1-Palmitoyl-2-oleoyl-*sn*-glycero-3-phosphoethanolamine (POPE) and 25% 1-Palmitoyl-2-oleoyl-sn-glycero-3-(phospho-rac-(1-glycerol)) (POPG) membrane. We used these percentages of lipid species in order to mimic the real membrane environment in which YidC is found. The full system was solvated with TIP3P water molecules (41) and 150 mM NaCl. The system consists of ~175 000 atoms in a triclinic box (~11×11×14 nm). The titratable groups of the protein were protonated according to their standard protonation states at pH 7. The preparation of the structure was done using the CHARMM-GUI webserver (42).

The initial, multi-step equilibration of the system, with a gradual release of restrains acting on protein atoms, was conducted using the scripts provided by the CHARMM-GUI. Subsequently, the system was equilibrated for 30 ns, without any restraints, prior to production runs.

#### Simulation details

All simulations were performed using the GROMACS 2020.3 simulation package (43) and the CHARMM36m force field (44). Ten independent simulations with different initial velocities were carried out cumulating 4 μs. All bonds were constrained using a fourth order LINCS algorithm (45), allowing for a 2 fs timestep. The short range interactions were cut-off beyond a distance of 1.2 nm. The long range electrostatic interactions were treated with PME (46) with a 1.2 nm real space cutoff. The systems were kept at a temperature of 303.15 K and a pressure of 1 bar, using a Nosé-Hoover thermostat (47, 48) and a Parrinello-Rahman barostat (49), respectively. Periodic boundary conditions were applied and snapshots were saved every 50 ps.

## Data Availability

Crystallographic data and coordinates were deposited in Protein Data Bank with accession number 6Y86. All remaining data are contained within this article.

## Supporting Information

This article contains supporting information.

## Acknowledgments

The PXI beamline scientists from Swiss Light Source, Villigen, Switzerland

## Author contributions

X.L and R.A.K. conceptualization and experimentation. K.J.N structure determination and Validation. I.M.I and A.C molecular dynamics simulations. M.J.S and A.J.M.D functional characterization. X.L, I.M.I and A.J.M.D making all figures. X.L writing original draft with inputs from all co-authors.

