## Supplementary Table 1-2 and Fig 1-4 for "The role of the first transmembrane helix in bacterial YidC: Insights from the crystal structure and molecular dynamics simulations"

1. Photon Science Division, Paul Scherrer Institute, Forschungstrasse 111, 5232 Villigen PSI, Switzerland
2. Department of Biochemistry, University of Zurich, Winterthurerstrasse 190, CH-8057, Zurich, Switzerland
3. Department of Molecular Microbiology, Groningen Biomolecular Sciences and Biotechnology Institute, Nijenborgh 7, 9727 AG Groningen, The Netherlands
   Current affiliation: Bioceros B.V. (Member of Polpharma Biologics), Yalelaan 46, 3584CM Utrecht, The Netherlands
4. Department of Molecular Microbiology, Groningen Biomolecular Sciences and Biotechnology Institute, Nijenborgh 7, 9727 AG Groningen, The Netherlands
5. Laboratory of Biomolecular Research, Division of Biology and Chemistry, Paul Scherrer Institute, Forschungstrasse 111, 5232 Villigen PSI, Switzerland.

a: contributed equally

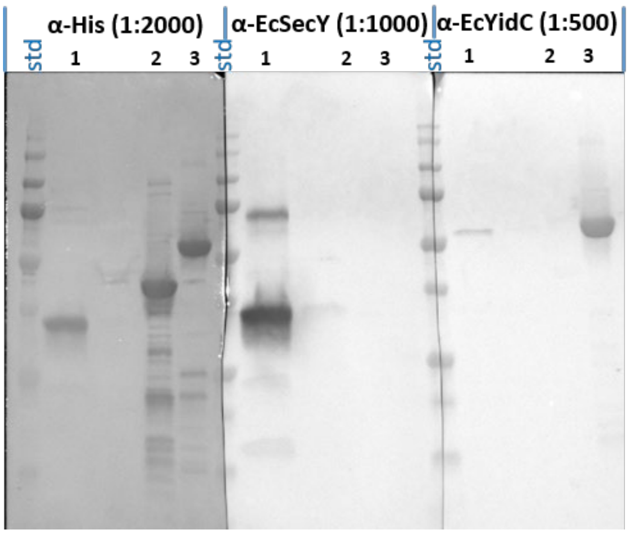


Figure 1: Interaction of EcSecY and EcYidC. Purified EcSecY, EcYidC and TmYidC all containing a hexa-his tag run on Bio-Rad anyKd gel and then blotted on nitrocellulose membrane. Three primary antibodies wereused to detect his-tagged protein (anti penta-His. 1:2000), EcSecY (anti-SecY rabbit,

SN810-2, 1:5000) and EcYidC (anti-EcYidC rabbit, self-made, 1:500). The secondary antibody for penta-His is goat-anti-mouse (1:4000) conjugated with Alkaline-Phoshatase and the secondary antibody for EcSecY and EcYidC is donkey anti-rabbit (1:4000) conjugated with alkaline phosphatase.  The blot was developed 30 sec for colorimetric detection.

```
T.maritima : MVLRKVVAIL-LAILPIFLFAVEPI : 24
E.coli     : MDSQRNLLVIALLFVSFMIWQAWEQ : 25
S.enterica : MDSQRNLLVIALLFVSFMIWQAWEQ : 25
K.pneumoni : MDSQRNLLIIALLFVSFMIWQAWEQ : 25
L.pneumoph : MDIRRIVLYMALALIGLSLWNAWQI : 25
Y.pestis   : MDSQRNLLLIALLFVSFMIWQAWQV : 25
H.influenz : MDSRRSLLVLALIFISFLVYQQWQL : 25
N.meningit : MDFKRLTAFFAIALVIMIGWEKMFP : 25
```

Figure 2: TM1 sequence alignment of different Gram-negative bacteria using ClustalX (1).

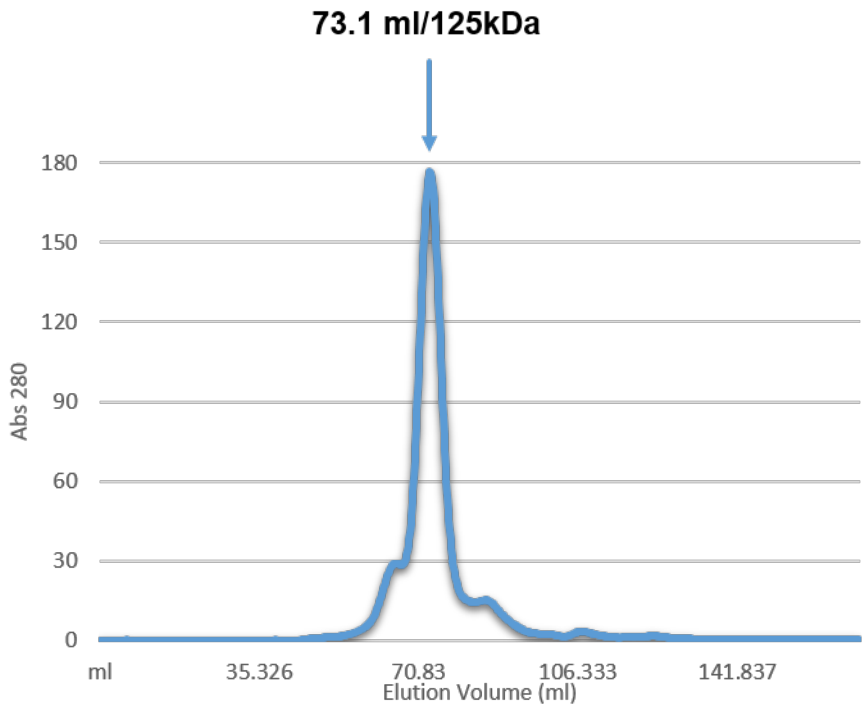


Figure 3: Gel filtration elution profile of purified TmYidC in the presence of 0.05%DDM. Purified TmYidC loaded on a Superdex 200 16 60 column using 50mM Hepes, pH 7.5, 150mM and 0.05%DDM buffer. The arrow pointed the elution volume of TmYidC and its molecular weight calculated based on the elution volume using the Bio-Rad standard running at the same condition.

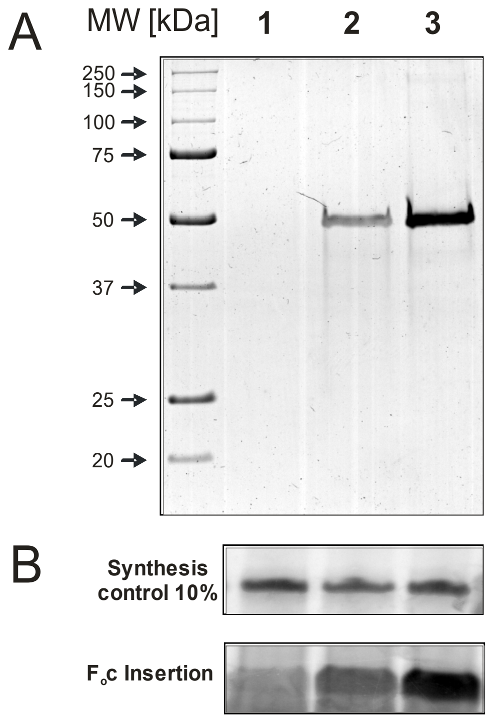


Figure 4**:** *Thermotoga maritima* YidC mediates the membrane insertion of in vitro synthesized *E. coli* $F_O$c. TmYidC was overexpressed in *E. coli* SF100 and affinity purified as described . (A) The purified protein was reconstituted in *E. coli* total lipids at different protein to lipid ratios and analyzed by SDS PAGE. Lane 1: no protein; lane 2: protein/lipid ratio 0.08 (wt/wt) , lane 3: protein/lipid ration 0.16 (wt/wt). Proteoliposomes were tested for the membrane insertion of in vitro synthesized $^{35}$S labeled *E. coli* $F_O$c. The total $F_O$c synthesis was equal in the presence of different proteoliposomes (B, upper panel), while the amount of membrane inserted $F_O$c (lower panel) increased with the TmYidC concentration in the reaction mixture.

**Strains and plasmids**

The sequences of all primers used in this study are listed in Table 1. All strains and plasmids are summarized in Table 2. In order to express TmYidC and EcYidC/TmYidC hybrid genes in *E. coli* cells, genes were cloned into the *E. coli* expression vector pTrc99A (Table 2). Constructs were verified by DNA sequencing. For expression of Tm_YidC with a $His_6$ tag attached in frame to its 3’ end, the gene was amplified from genomic DNA of *T. maritima* MSB8 (generous gift from S.V. Albers) (2) using the primer pair FoTmaNcoI/ReTmHisXbaI (Table 1). The PCR product was cleaved with NcoI and XbaI and ligated into the corresponding sites of the vector pTrc99A, yielding pTrcTmHis (Table 2).

The Ec and Tm_YidC hybrids were constructed by overlapping PCR. The first (5') 186, 963 and 1038 bps of *yidc* were amplified from genomic DNA of *E. coli* DH5α using the forward primer FoYidCXbaI and either ReYidC61Tma396, ReYidC321Tma225 or ReYidC346Tma198 as reverse primer (Table 1). Subsequently, DNA fragments consisting of the last (3') 1188, 675 and 594 bps of *tm1461* were amplified from genomic DNA of *T. maritima* MSB8 using either FoYidC61Tm396, FoYidC321Tm225 or FoYidC346Tm198 as forward and ReTmHisHindIII as reverse primer. The fragments 186 and 1188, 963 and 675, and 1038 and 594 were fused by overlapping PCR using the combined DNA fragments as template and the primer pair FoYidCXbaI/ReTmHisHindIII. The PCR products were cleaved with XbaI and HindIII and ligated into the corresponding sites of pTrc99A, yielding pTrce61-t396His, pTrce321-t225His and pTrce346-t198His.

Table 1. Primers.

| Primer | Sequence (5'→3') |
|---|---|
| FoTmNcoI | GCTGCCATGGTCTTGAGAAAAGTTGTAGC |
| ReTmHisXbaI | GCGTCTAGATCAGTGATGGTGATGGTGATGTGCCTTTTTCGGAAGACCAAGAAGTTCTCTC |
| FoYidCXbaI | GCTCTAGATGGATTCGCAACGC |
| ReTmHisHindIII | CACGAAGCTTTCAGTGATGGTGATGGTGATGTGCC |
| FoYidC61Tm396 | GTGGCCAGGGGAAACTGATCTCGGGCATACTCAAAGACTTTTACACCCTT |
| ReYidC61Tm396 | CGAGATCAGTTTCCCCTGG |
| FoYidC321Tm225 | GACAAAATGGCAGCTGTTGCTCCAGGATTCAACAAGTGGTA |
| ReYidC321Tm225 | TACCACTTGTTGAATCCTGGAGCAACAGCTGCCATTTTGTC |
| FoYidC346Tm198 | TGTTCAAACTGCTGAAATGGTTCGGATGGGCGATCATGCT |
| ReYidC346Tm198 | AGCATGATCGCCCATCCGAACCATTTCAGCAGTTTGAACA |

Table 2. Strains and plasmids.

| Plasmids or strain | Relevant properties | Reference |
|---|---|---|
| Plasmids | | |
| pTrc99A | Expression vector for *E. coli* based on pKK233-2, carries a hybrid *trp/lac* promoter and the multiple cloning site of pUC18; $Ap^R$ | (3) |
| pTrcyidC | Like pTrc99A, contains *E. coli yidC* | (4) |
| pTrcTmaHis | Like pTrc99A, contains *T. maritima tm1461* with a $His_6$ tag at 3' end | This study |
| pTrce61-t396His | Like pTrc99A, contains the 3' 183 bps of *E. coli yidC* fused in frame to the 5' 1188 bps of *T. maritima tm1461* with a $His_6$ tag at 3' end | This study |
| *E. coli* strains | | |
| DH5α | *supE44 hsdR14 recA1 endA1 gyrA96 thi-1 relA1 ΔlacU169* (Φ80*lacZ* ΔM15); K12 derivative | (5) |
| SF100 | $F^-$ *ΔlacX74 galK thi rpsL (strA) ΔphoA(pvuII) ΔompT* | (6) |
| NN100 | SF100, *lpp D*(*uncB-C*) *zid*::*Tn10*) | (7) |
| FTL10 | Δ*yidC attB*::(*araC*$^+$ $P_{BAD}$ *yidC*$^+$); $Kan^R$ | (8) |

**In vivo complementation**

In *E. coli* strain FTL10 (Table 2) the *yidC* gene is under control of the araBAD promotor/operator. Because YidC is an essential protein in *E. coli*, cell growth of the FTL10 strain is strictly dependent on the presence of arabinose in the medium. The *E. coli* FTL10 was transformed with plasmids pTrc99A (ΔYidC), pTrcyidC (wild-type), pTrcTmHis (*T. maritima* TM1461), pTrce61-t396His (YidC61), pTrce321-t225His (YidC321) or pTrce346-t198His (YidC346). Cells were streaked out on LB agar plates containing 100 μg/ml ampicillin and glucose ($LB^{Amp}$ agar), and incubated for at least 16 h at 37 ºC. To examine cell growth in liquid culture, at least three single colonies from each of the above-mentioned strains were picked and overnight culture grown at 37 °C in LB medium containing 100 µg/ml ampicillin and 0.5% (wt/vol) arabinose. Next morning, precultures were diluted 100-fold in prewarmed LB medium containing 0.5% (wt/vol) glucose and 50 µM IPTG. The optical density at a wavelength of 660 nm ($OD_{660}$) was measured at different time points until cessation of cell growth.

**Expression and Purification of YidC of *Thermotoga maritima***

The plasmid pTrcTmaHis (Table 2) was transformed into competent *E. coli* NN100 cells using the heat-shock method and plated on $LB^{Amp}$ agar. Single colony appeared on $LB^{Amp}$ agar incubated at 37°C overnight was inoculated into 2 ml $LB^{Amp}$ medium and shaken at 37°C for 7h-8h at 180rpm/min. The 2ml $LB^{Amp}$ culture was then transferred into 200ml $2XTY^{Amp}$ and shaken at 37°C for 16-17h at 180rpm/min. The 15ml overnight culture was then inoculated into 1L $LB^{Amp}$ for 12L large scale expression. The large bacterial cultures were shaken at 30°C with 200rpm/min. 0.5mM IPTG was added into the culture when the OD600 reached 0.8-1.0. Cells were harvested after 2h after IPTG induction. Harvested cells were either used directly or frozen at -80°C.

Cells from the 12L cultures were resuspended in cold lysis buffer (50 mM Na-phosphate (pH 8.0), 300 mM NaCl, 10% glycerol, and 10 mM imidazole] by gentle shaking. After cell disruption using a French press (Emulsiflex C3) the lysate was centrifuged for 30 min at 10,000 x *g*. Membranes were pelleted by centrifugation of the supernatant at 100,000 x *g* for 1 h and were resuspended in lysis buffer, potter homogenized, and the protein solubilized with 1% (wt/vol) dodecyl-maltoside (DDM). The supernatant obtained after ultracentrifugation was subjected to a cobalt affinity chromatography carried out using batch method. In average, the solubilized 2.5 g membranes were incubated with 1ml cobalt beads at 4°C with a rotator for at least 1h. The beads were washed with 20 ml of lysis buffer containing 0.05% DDM, followed by 10 ml buffer containing 20 mM imidazole and subsequently with 10 ml buffer containing 40 mM imidazole. The protein was eluted with 5x1ml elution buffer containing 300 mM imidazole. The elution fractions 2 to 4 were pooled and concentrated to 1 ml, desalted on a NAP-10 column, using desalting buffer [(50 mM TrisHCl pH 8.5, 100 mM NaCl, 10% glycerol, and 0.12% decyl maltoside (DM)] and concentrated again to >10 mg/ml by the nanodrop method.

**Reconstitution and in vitro transcription, translation and membrane insertion**

Purified *E. coli* $F_oc$ and TmYidC were reconstituted into *E. coli* phospholipids (Avanti Polar Lipids, Alabaster, NY) using Bio-beads SM-2 (Bio-Rad) as described {van der Laan, 2001 #594}. Both proteins were reconstituted at protein/lipid ratios of 0.08 and 0.16 (wt/wt).

For in vitro synthesis of *E. coli* $F_oc$ using pET20batpE as DNA template, a cell lysate was prepared essentially as in {Saller, 2009 #412}. Coupled transcription, translation and membrane insertion reactions were performed as described previously {Jewett, 2004 #638} with DTT present at a final concentration of 4 mM. Each reaction was started by addition of T7 polymerase from Fermentas and the Easytag express protein labeling mix (Perkin Elmer). In vitro synthesis and insertion were carried out for 30 min at 37°C in the presence of proteoliposomes. A 10% synthesis control was collected, and the remainder of the mixture was subjected to proteinase K treatment (2 mg/ml, 30 min, 37°C) in order to remove non-inserted proteins. All consecutive steps were performed as described previously {van der Does, 2003 #639}.

**Crystallization and Data collection**

Crystallization trials were set up in sitting drops using a Phoenix liquid handling system (Art Robbins Instruments) and different, commercially available, screens (Nextal). Initial crystals diffracting to approximately 25 Å were obtained from several conditions containing PEG3350 as precipitant. A homemade screen with varying pH and PEG3350 concentration was used for optimization and batch-to-batch variation. Detergents, lipids, heavy atoms and small molecules were tested as additives. Crystals diffracting to high resolution were obtained after 2-7 days at 20°C from sitting drops mixed in a 1:1 ratio with crystallization solution 0.1M Bis-Tris, pH5.5, 0.2M di-ammonium hydrogen citrate with 24% PEG 3350.

For data collection, crystals were mounted on cryoloops directly from crystallization drops and frozen in a cold nitrogen gas stream. Data were recorded at 100 K using a rotation angle of 1.0° per image on a MarCCD detector at the protein beamline X06SA, Swiss Light Source (SLS), Paul Scherrer Institut, Villigen. Datasets were processed with the XDS program (4).
